# The KDM6 histone demethylase inhibitor GSK-J4 induces metal and stress responses in multiple myeloma cells

**DOI:** 10.1101/2024.12.28.630531

**Authors:** Adam P Cribbs, Edward S Hookway, Clarence Yapp, Ka-Hing Che, James E Dunford, Karina Gutheridge, Federica Lari, Graham Wells, Martin Philpott, Peter Cain, Deborah Brotherton, Charlotte Palmer, Wolfgang Maret, Jude Fitzgibbons, John C. Christianson, Udo Oppermann

## Abstract

Thioneins are cysteine-rich apoproteins that regulate divalent metal homeostasis by virtue of their metal-chelation properties resulting in the ligand-bound metallothionein state. Previous studies show transient upregulation of the metallothionein (MT) gene cluster as part of a complex transcriptional response to a class of histone demethylase tool compounds targeting human Fe^2+^ dependent ketoglutarate oxygenases KDM6A (UTX) and KDM6B (JmjD3). The prototypic bioactive KDM6 inhibitor GSK-J4 induces apoptotic cell death in multiple myeloma cells and corresponding transcriptomic profiles are dominated by metal and metabolic stress response signatures, also observed in primary human myeloma cells. Here we investigate the hypothesis that metal-chelation by GSK-J4 provides the means for transport and intracellular release of Zn^2+^ leading to a metallothionein transcriptomic response signature. Live cell imaging of myeloma cells shows transient increases in intracellular free Zn^2+^ concentrations when exposed to GSK-J4, consistent with a model of inhibitor-mediated metal transport. Comparisons of GSK-J4 and ZnSO_4_ treatments in the presence or absence of metal chelators show that both treatment conditions induce different transcription factor repertoires with an overlapping MTF1 transcriptional regulation responsible for metallothionein and metal ion transport regulation. The data provide a possible explanation for the observed metal response upon GSK-J4 inhibition however the relationship with the pro-apoptotic mechanism in myeloma cells requires further investigation.

## INTRODUCTION

Metal ions derived from iron, copper, zinc, cobalt, chromium, manganese, magnesium, calcium, molybdenum, and non-metal elements such as selenium play fundamental roles in biochemistry. They participate in processes such as transport and utilisation of oxygen, in metabolic pathways as well as in transcriptional regulation and accordingly are required to maintain homeostasis in virtually all aspects of cellular physiology. In humans, both metal deficiencies and metal excess are causative in rare disorders or are involved in cancer or development in a metal-specific manner, including for example abnormal zinc levels leading to growth retardation or altered immune functions [1, 2]. Redox-inert metal ions such as Zn^2+^ are co-factors for approximately 10% of human proteins, are required for catalytic activity and structural integrity of proteins, or constitute in the case of calcium, and more recently also postulated for zinc, second messengers in cellular signalling processes [3].

Levels of free metal ions in cellular systems are tightly regulated and range from atto- to micromolar concentrations, dependent on the metal ion, while total concentrations are often significantly higher due to exquisite proteinaceous buffering systems present within the cell. These buffering and membrane transport systems provide the homeostatic control that maintains essential transition metal ions at characteristic cellular concentrations. A paradigm for biological metal buffering are metallothioneins, isolated in 1950’s as a cadmium binding protein from horse kidney by Vallee and Margoshes [4]. In humans, metallothioneins constitute a cluster of 14 genes, located on chromosome 16q13 [5], encoding structurally similar, 60-odd amino acid residue, cysteine-rich intracellular metal-binding proteins [6]. Metal ligands include zinc and copper, but also others such as cadmium, mercury, platinum as well as silver, providing protection to cells and tissues against heavy metal toxicity [7]. Whilst this toxicological defense mechanism has been postulated early on, more recently metallothioneins have been implicated in normal physiological processes such as hematopoietic cell proliferation and differentiation, and in disease, such as cancers, through their ability to control zinc levels and maintain redox homeostasis [8–11].

Transcription of metallothionein genes, as well as other genes involved in zinc metabolism such as the zinc exporter SLC30A1 are regulated by the metal-response element (MRE) – binding transcription factor 1 (MTF1) [12]. Upon metal exposure or cellular changes in the availability of metal ions, MTF1 binds zinc and other metal ions in the cytosol, initiating nuclear translocation and chromatin binding of MTF1 to metal response elements (MRE) found in promoters of metallothioneins and other target genes.

Metal chelation is an important therapeutic principle in treatment of metal intoxication and in several metabolic and fibrotic diseases, in anti-cancer or anti-infective therapies, where metalloenzymes are selected as drug targets. Inhibitor chemotypes for these metalloenzymes are substrate or co-substrate mimics containing metal chelation motifs. One such example is the prototypic cell-permeable inhibitor GSK-J4, which specifically inhibits the histone lysine demethylases KDM6A and KDM6B [13]. These enzymes constitute an important class of epigenetic regulators such as” readers, writers and erasers” of a histone code [14–16]. KDM6 enzymes play an important role in regulation of chromatin modifications and belong to the “eraser” group by specifically demethylating methylated histone 3 lysine27 (H3K27me1/2/3/) residues [17]. This lysine methylation constitutes the enzymatic product of the polycomb repressive complex 2 (PRC2) and plays a central role in cellular physiology [18, 19].

Considering the critical role of metal homeostasis in oncological processes and the inherent metal-chelating capabilities of GSK-J4, our study aims to investigate the effects of GSK-J4’s inhibition of KDM6A and KDM6B on metal ion regulation in myeloma cells. We hypothesize that GSK-J4 mediates its effects not only through epigenetic modifications but also by altering cellular metal landscapes, potentially impacting disease outcomes. This investigation aims to bridge the gap between metal homeostasis and epigenetic regulation, offering new insights into the therapeutic mechanisms of histone demethylase inhibitors in cancer treatment.

## RESULTS AND DISCUSSION

### A focused epigenetic compound library screen identifies the KDM6 inhibitor GSK-J4 as anti-proliferative agent across multiple myeloma cell lines

We compiled a focused library consisting of >120 validated and well-characterised epigenetic tool compounds to interrogate epigenetic biology, either in-house developed or by using commercially available sources **(SI DATA 1)**. We have successfully used this approach to investigate and dissect the roles of multiple epigenetic modulators in several areas including oncology, inflammation or stem cell biology [20–25]. Using a colorimetric resazurin-based metabolic viability read-out as proxy for anti-proliferative activity we now assessed activities of epigenetic tool compounds over 72 hrs against 4 multiple myeloma cell lines **(Figure 1A, SI DATA 1)**. The library screen clearly identified several epigenetic classes previously shown to inhibit myeloma cell proliferation such as inhibitors for BET bromodomains (eg BRD4) [26], histone deacetylases [27, 28], or methyltransferases [29], as well as other known metabolic targets such as proteasome or tRNA synthetase inhibitors [30], which we included as positive controls. In addition we here show that the KDM6 inhibitor series GSK-J4, including its derivative KDOBA67 [22] is highly active across all myeloma cell lines tested. The GSK-J4 series **(Figure 1B)** was developed as a selective, cell-permeable, bioactive tool compound set to interrogate histone demethylase KDM6 biology by using a structure-guided approach [13]. Apart from LSD1 (KDM1), histone demethylases belong to a family of evolutionarily conserved 2-oxoglutarate and Fe^2+^ dependent oxygenases [31, 32]. GSK-J1, the intracellularly active molecule, has an IC_50_ of 60 nM for KDM6B in an *in vitro* demethylase assay and binds with its amino-propyl moiety in the 2-oxoglutarate pocket of KDM6, whilst the pyridyl-pyrimidine biaryl of GSK-J1 makes a bidentate interaction with the catalytic metal **(Figure 1C)**. This metal-chelating binding mode is critical for demethylase inhibition. The pyridine regio-isomer GSK-J2 cannot form such an interaction and hence displays considerably weaker KDM6 inhibition (IC_50_ > 100 μM) making it an ideal negative control compound to investigate the functional role of KDM6 enzymes. To overcome the issue of cell permeability often encountered with carboxyl-containing molecules, ethyl esters of GSK-J1 and GSK-J2 (GSK-J4 and GSK-J5, respectively) were synthesized **(Figure 1B)**. This pro-drug approach provides cell permeability and on-target activity after intracellular cleavage of the carboxyl-ester of GSK-J4 and GSK-J5 which is mediated by unspecific intracellular hydrolases. Accordingly, GSK-J5 does not show anti-proliferative activity in our cell-based screen **(Figure 1A)**.

**Figure 1:**
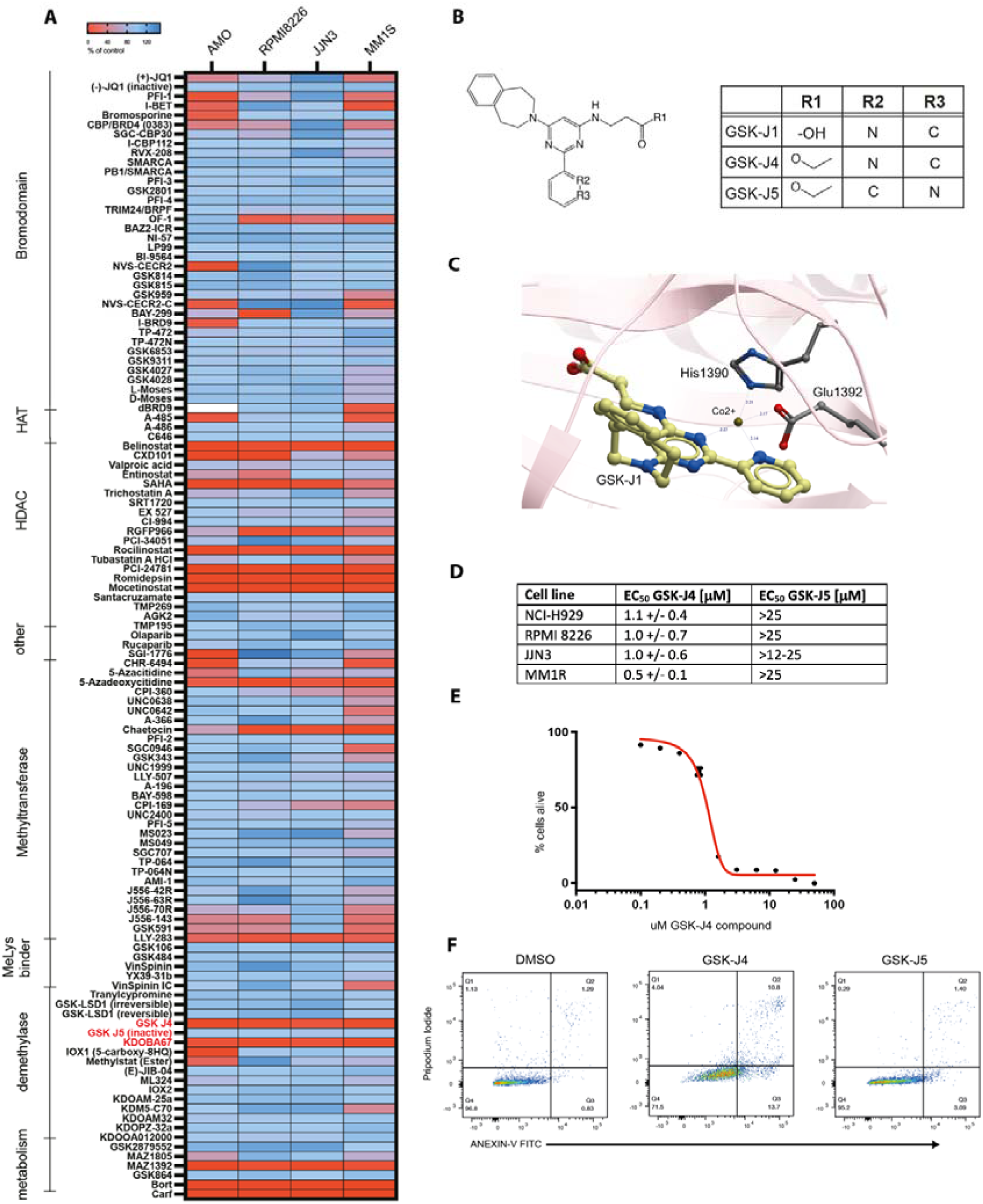
GSK-J4 is a bioactive tool compound for inhibiting KDM6 enzymes and induces a metal response signature and apoptosis in myeloma cells. **(A)** Heatmap reporting as % change over solvent control (DMSO) in the viability of multiple myeloma cell lines (AMO1, JJN3, RPMI8226, MM1S), screened against a focused library of epigenetic tool compounds (SI Data 1) as measured using PrestoBlue assay. **(B)** Structures of GSK-J4 and analogues used in this study. **(C)** Binding mode and chelation of active site metal in human KDM6B by GSK-J1 (presented as sticks, PDB code 4ASK). Sphere depicts position of the Co2+ metal used in the crystallisation experiments, amino acid residues of KDM6B (His1390, Glu1392) involved in metal coordination are depicted. **(D)** EC_50_ values for anti-proliferative effects measured in multiple myeloma cell lines treated with GSK-J4 or GSK-J5. **(E)** Dose response curve for GSK-J4 in JJN3 cells (EC_50_ = 1.1 μM). **(F)** Flow cytometry data for JJN3 cells and induction of cell death upon 72 hours of DMSO, GSK-J4 and GSK-J5 treatment (2 μM).

### The KDM6 inhibitor GSK-J4 has potent anti-proliferative activity in multiple myeloma cell lines

We determined EC_50_ values for several myeloma cell lines showing low-to submicromolar EC_50_ values (**Figure 1D**), with an EC_50_ of 1.0 +/-0.6 μM for GSK-J4 in the JJN3 multiple myeloma cell line **(Figure 1D, 1E)**. Consistent with a metabolic impairment indicating anti-proliferative effects upon inhibitor treatment we observed a reduction of DNA replication as measured by BrDU incorporation, showing a reduction of the proportion of cells in S and G2/M and an increase in sub-G1 cell cycle phases in JJN-3 cells **(SI Figure 1)**. Furthermore, JJN3 cells treated with GSK-J4 for 72 hrs exhibited increased expression of apoptosis marker ANNEXIN-V as well as propidium iodide staining by flow cytometry, indicating the activation of a pro-apoptotic mechanism leading to increased cell death **(Figure 1F, Supplementary Figure 1, Supplementary Figure 2, Supplementary Figure 3 and Supplementary Figure 4)**.

### Transient upregulation of metallothionein genes in myeloma cell lines and patient bone marrow is a pervasive transcriptional signature of GSK-J4 treatment

To better understand the GSK-J4 mediated response, we performed transcriptomic analyses using several different methodologies. Firstly, microarray studies of JJN3 cells treated with >EC_50_ (2 mM) GSK-J4 over 6 or 24 hr exhibited a strong upregulation of metallothionein genes including *MT1E, MT1F, MT1G, MT1H, MT1X* and *MT2A* while the inactive analogue GSK-J5, did not show any significant transcriptional changes **(Figure 2A)**. Metallothionein up-regulation correlated with increased expression of the zinc transporter *SLC30A1*, suggesting a potential biological relationship between KDM6 inhibition and zinc homeostasis. This effect was also observed in RPMI myeloma cells. The dominant transcriptional metallothionein induction is also observed at sub-EC_50_ (0.5 mM) at 24 hr treatment. This metallothionein response was distinct from that produced by bortezomib – a proteasome inhibitor known to disrupt protein homeostasis in myeloma. In addition to this metal response, a signature of upregulated ATF4 target genes was found in both the GSK-J4 and bortezomib treated cells, reflecting induction of the integrated stress response (ISR) by these compounds **(Figure 2A)** [23, 33]. By re-analysing previous datasets comprising a wide array of cell-based systems from our and other laboratories [23, 25, 34–36] we indeed detect a significant induction of metallothionein genes upon GSK-J4 exposure **(Supplementary Figure 5)**, suggesting metallothionein induction as a generic feature of the transcriptional response upon GSK-J4 treatment. We next performed bulk RNA sequencing of JJN3 cells treated with GSK-J4 in a time series for 3hr, 6hr and 24hr **(Figure 2B-D)**. These data revealed a transcriptional signature that consisted of over 2060 genes showing a significant log2-fold change across all timepoints. From the upregulated gene set, an enriched MTF1 transcription factor motif was identified **(Figure 2E)**, further confirming the presence of a zinc response following GSK-J4 treatment. Similarly, an enrichment of genes associated with the ATF4 transcription factor motif was also observed **(Figure 2E)**. In line with the observed data, Gene Ontology analysis of the transcriptional data at 24 hrs confirmed stress responses to Zn^2+^, regulation of the execution phase of apoptosis and an endoplasmic reticulum unfolded protein response **(Figure 2F)**. The metallothionein response was detectable at 3 hrs, was maximal at 6 hrs but began to decline after 24 hrs **(Figure 2G)**. These trends were confirmed by using qPCR of the MT1X gene as surrogate marker, which showed reduced expression after treatment with GSK-J4 for 12 hrs **(Figure 2H)**.

**Figure 2:**
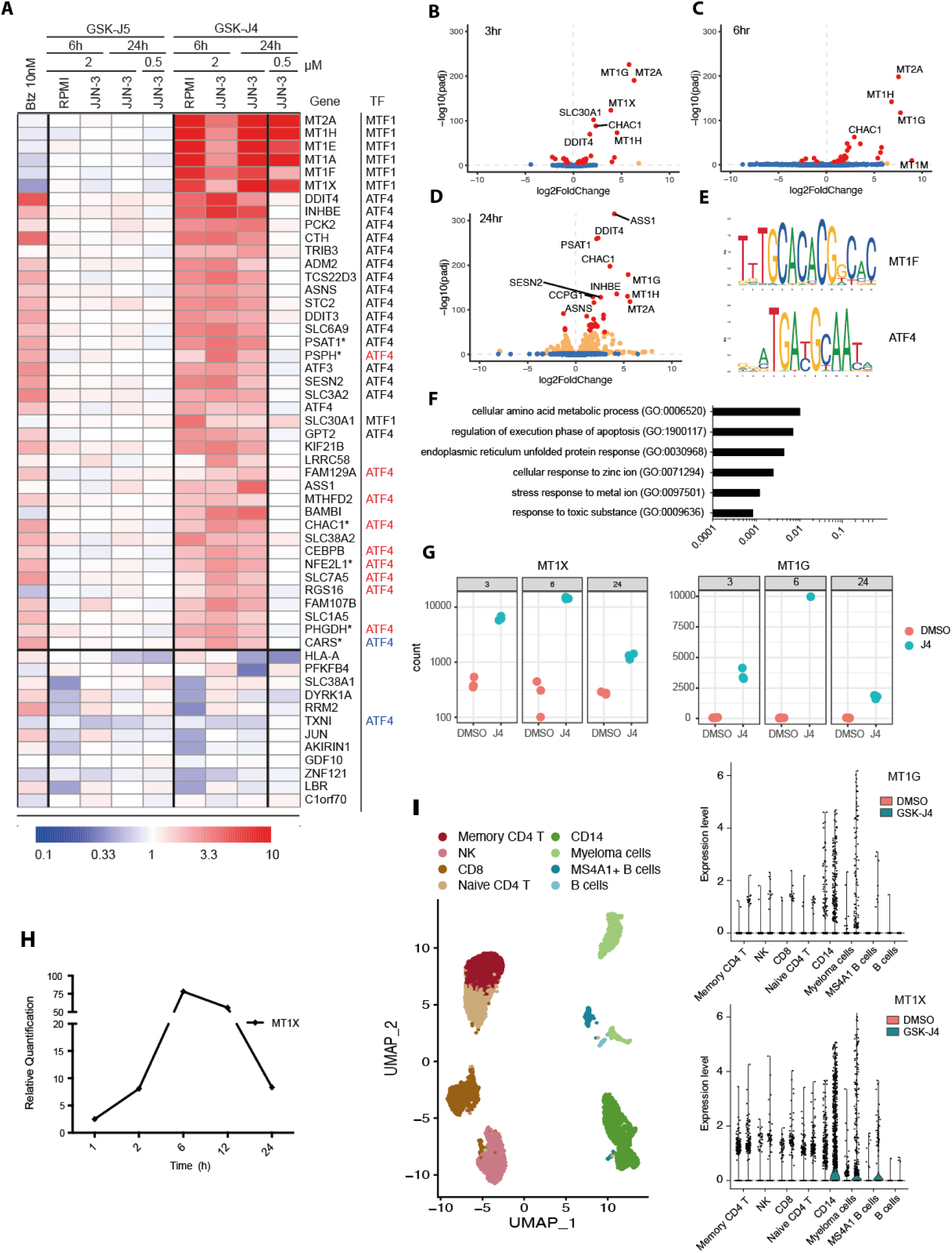
GSK-J4 induces an ATF4- and MTF1-dependent transcriptional response. **(A)** A heatmap displaying the differentially regulated genes identified by microarray analysis in JJN3 and RPMI cell lines treated with various concentrations of DMSO (0.1%) or GSK-J4 (2 μM and 0.5 μM) concentration. The map highlights the known associations with the transcription factors ATF4 or MTF1, and includes a contrast with JJN3 cells treated with the proteosome inhibitor Bortezomib (10 nM) **(B-D).** Volcano plots showing selected differential gene expression responses from RNA-seq data for **(B)** 3 hrs, **(C)** 6 hrs and **(D)** 24 hrs following GSK-J4 treatment (2 μM). **(E)** Motif enrichment analysis showing the top two enriched motifs at the 24 hr timepoint. **(F)** GO analysis showing the top enriched ontology terms at the 24 hr timepoint. **(G)** Scatter plots showing the expression of MT1X and MT1G following treatment of JJN3 cells with either 3 hrs, 6 hrs or 24 hrs of DMSO or GSK-J4 treatment (2 μM). **(H)** RT-PCR analysis of MT1X gene expression of JJN3 cells treated with GSK-J4 for 1, 3, 6, 12 and 24 hrs. **(I)** UMAP of single-cell sequencing of human bone marrow samples (newly diagnosed myeloma), depicting the major cell type clusters identified. The top right panel shows the expression of MT1X treated with either DMSO (laft bar) or GSK-J4 (right bar; 2 μM). The bottom right panel shows the expression of MT1G treated with either DMSO (left bar) or GSK-J4 (right bar; 2 μM).

To investigate whether GSK-J4 produced a comparable response in myeloma patient derived cells, we exposed a myeloma bone marrow aspirate to GSK-J4 for 24 hrs and measured the transcriptional response by single cell sequencing **(Figure 2I, Supplementary Figure 6)**. In addition to myeloma cells, we also identified clusters of myeloid and lymphoid cells (B-cells, T-cells, NK-cells). Expression analysis reveals that GSK-J4 treatment induces upregulation of metallothionein genes MT1X and MT1G in the myeloma and myeloid and to some extent in the lymphoid cell clusters. The data confirm that the GSK-J4 induction of metallothionein genes, as observed in previous studies in peripheral blood-derived immune cells occurs also in bone marrow immune populations, including myeloma cells. The role of metallothioneins in the immune system is largely underexplored but a role in immunity has been postulated previously [37].

### Modulation of intra- and extracellular Zn^2+^ levels regulate metallothionein gene expression

We next investigated whether the upregulated metallothionein response to GSK-J4 originated from changes in levels of intracellular or extracellular Zn^2+^. JJN3 cells exposed to GSK-J4 (1hr and 3hr) were treated with either TPEN or EDTA, Zn^2+^ chelators that are cell-permeable and non-permeable, respectively [38, 39] and the transcriptional metallothionein response was monitored **(Figure 3A-B)**. A similar response pattern was observed in AMO1 and MM1S cell lines, measured at the 3hr timepoint **(Figure 3-D)**. Both TPEN and EDTA significantly reduced levels of MT1X in the DMSO treated control as measured by qPCR, indicating a very tight control of the MT gene locus by Zn^2+^ levels. Importantly, the GSK-J4 induced metallothionein response at 3hrs (about 10-fold induction eg in JJN3, about 100-fold in AMO1 cells) is completely abrogated by the Zn^2+^ chelator (**Figure 3B-D**), suggesting that metal levels sensed through MTF1 may be the primary stimulus for the observed transcriptional metallothionein response observed with the KDM6 inhibitor. Bioavailable zinc in the form of ZnSO_4_ yields a 10-fold induction by 1 hr, upregulating to nearly 100-fold by 3 hrs (**Figure 3A, B**). This strong upregulation can only be partially blunted by the addition of chelating agents, indicating a dominant and tight MTF1 regulation of the transcriptional metal response. While MT2A, MT1X, MT1E and MT1G are clearly upregulated at the 3-hr timepoint, the magnitude of metallothionein induction differs between GSK-J4 and ZnSO_4_ treatments which may indicate a concentration-dependent effect of Zn^2+^ on the transcriptional response.

**Figure 3:**
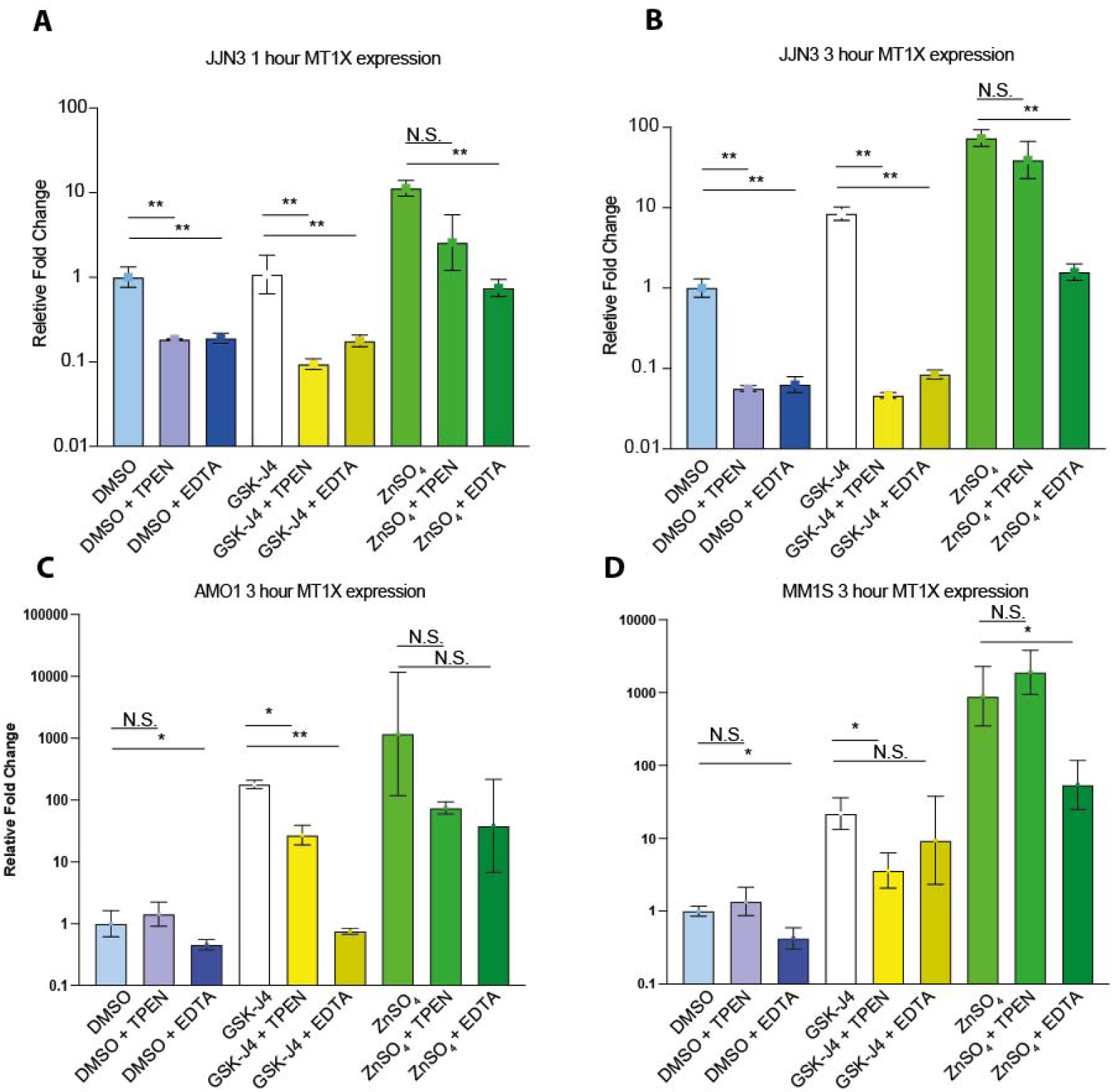
Modulation of intra- and extracellular Zn2+ levels regulates MT1X expression. – MT1X expression was monitored by qPCR at 1 hr **(A)** and 3 hr **(B)** timepoints after DMSO (0.1%), GSK-J4 (2 μM) or ZnSO4 (100 μM) exposure in the absence or presence of extracellular (EDTA; 1 mM) or intracellular (TPEN; 10 μM) Zn chelators. The experiment was then repeated at the 3hr timepoint in AMO1 **(C)** and MM1S **(D)** cells. P values were calculated using Kruskal-Wallis test followed by Dunn’s multiple companrrison. *P <0.05, ** P<0.01. Error bars show mean +/- SD.

### GSK-J4 treatment increases intracellular zinc levels in myeloma cells

The transcriptional response elicited by GSK-J4 treatment strongly indicates a role for divalent metal ions. Given the effect of chelators, we hypothesized that an increase in intracellular Zn^2+^ ions might be involved in the process. To investigate this, we used Fluozin-3 AM staining and live cell imaging by fluorescence microscopy to measure intracellular free Zn^2+^ concentrations within single JJN3 cells (**Figure 4**) [40]. Following GSK-J4 treatment, intracellular free Zn^2+^ levels transiently increased with a peak observed at 50 min of treatment (**Figure 4A**), followed by a decline in free Zn^2+^ levels. Importantly, exposure of myeloma cells to ZnSO_4_ (200 μM) does not reduce cell viability (**Figure 4B**) and we do not observe a shift in the EC_50_ of GSK-J4 when JJN3 cells are co-treated with 20μM ZnSO_4_ (**Figure 4C**). These data are consistent with a Zn^2+^ transport into cells directly mediated by GSK-J4, possibly through the pyridine-pyrimidine chelating element of GSK-J4. Such a model implies chelation of Zn^2+^ by GSK-J4 from the media, transport of the metal-inhibitor complex across the plasma membrane, and subsequent release of Zn^2+^ within the cell.

**Figure 4:**
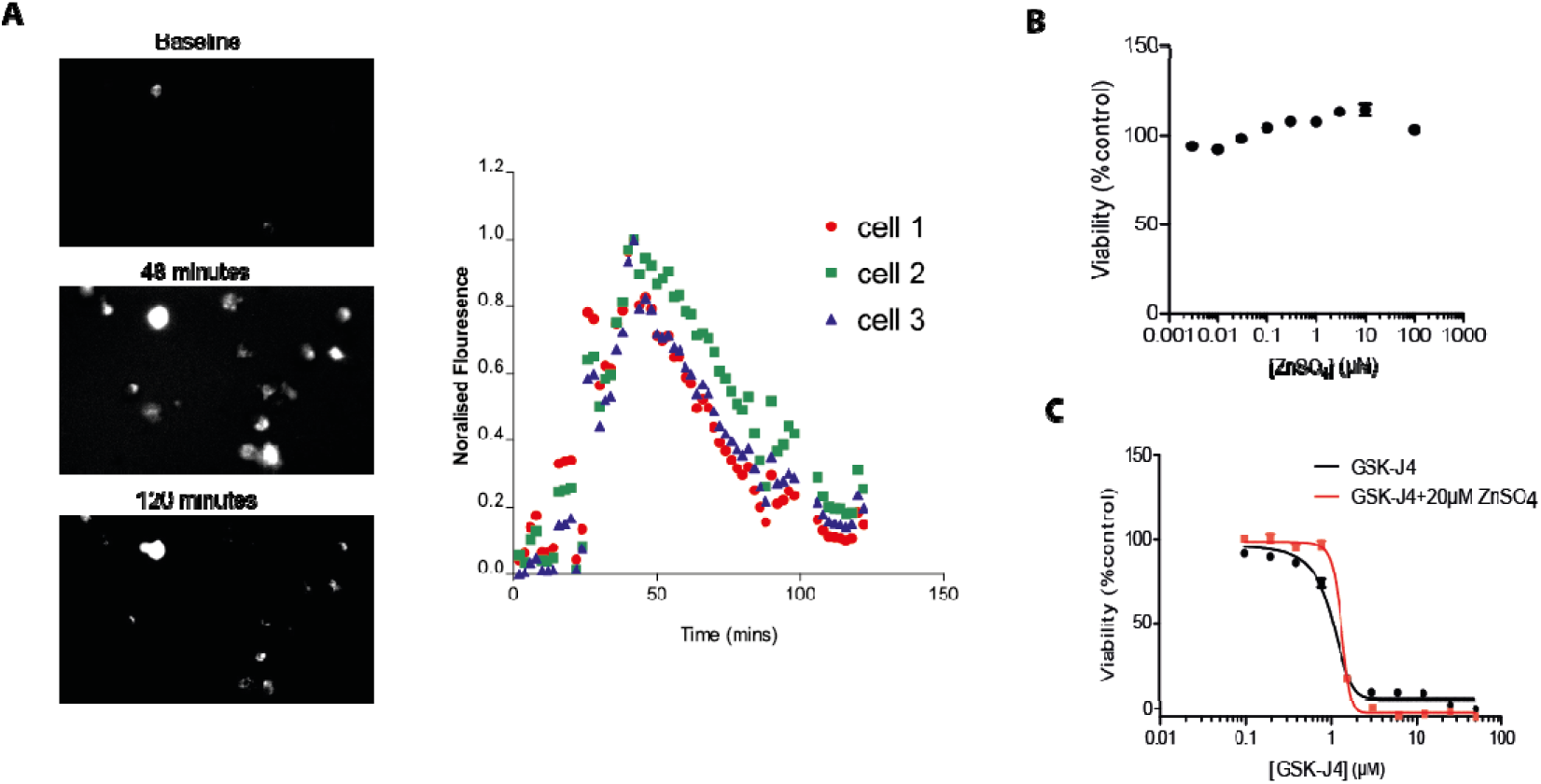
GSK-J4 treatment induces a transient surge in free intracellular Zn2+. **(A)** Left panel shows representative images of FluoZin-3 stained JJN3 cells at baseline, 48 mins and 120 mins following GSK-J4 treatment (2 μM). The right panel shows the relative fluorescence of FluoZin-3 in 3 representative cells over 120 mins of exposure with GSK-J4. **(B)** Dose response curve for ZnSO_4_ treatment in JJN3 cells. **(C)** Dose response curve for GSK-J4 treatment alone or co-treatment of GSK-J4 and ZnSO_4_ in JJN3 cells.

### GSK-J4 and ZnSO_4_ treatment share a common metallothionein and stress response but induce different transcriptional programs and intracellular pathways

Above we have shown that GSK-J4 induces a transient increase in intracellular zinc and induces cell death in JJN3 cells (**Figures 4A, 1F**) but that Zn^2+^ alone (through ZnSO_4_) does not induce cell death (**Figure 4C**). To identify gene signatures that may be related to induction of cell death by GSK-J4, it was necessary to try and uncouple the temporal dynamics of the Zn^2+^ response from the overall GSK-J4 response. We monitored transcriptional changes in JJN3 cells treated with either ZnSO_4_ or GSK-J4 over a 24-hour time course. Using a cut-off value of 50 in base mean expression across the different conditions, both ZnSO_4_ and GSK-J4 resulted in changes in gene expression over the 24-hr time scale, with 83 differentially regulated genes in the ZnSO_4_ treatment and 124 differentially expressed genes with GSK-J4 **(Figures 5A, 5B)**. Apart from the metallothionein response genes, bulk RNA-seq data revealed few upregulated genes that overlapped between GSK-J4 and ZnSO_4_ conditions. Both treatments regulate genes involved in intrinsic apoptotic signalling pathways and nutrient responses such as HMOX, SESN2, BBC3, ATF3 and DDIT4 **(Figures 5C, 5D)**. Zn^2+^ specific responses are dominated by induction of heat shock proteins and chaperones such as HSPA6, DNAJA4 and DNAJB1, while KDM6 inhibitor (GSK-J4) specific genes involve elements of the ISR and include INHBE, DDIT3, ATF4, HERPUD1 and ERN1 **(Figure 5E)**. Previous work using a MTF1 depletion approach, demonstrated the existence of a network of metal response factors mediating transcriptional responses to Zn2+ treatment, with MTF1 controlling a postulated hierarchical structure of Zn2+ sensors [41]. Transcription factors control expression of critical genes and thus may play a prominent role in eliciting the differences seen between GSK-J4 and ZnSO_4_ treatments. We next investigated the transcription factor repertoires that each treatment produced. Consistent with an ATF4 dominated response, GSK-J4 treatment leads to the upregulation of several transcription factor genes associated previously with the ISR. In particular, increases were observed for DDIT3, but also CEBPB. Upregulation of chromatin factors such as histone demethylase KDM4A and SS18L1 as part of the BAF complex may indicate an epigenetic component as part of the GSK-J4 response (**Figure 5F**). In contrast, few transcription factors were differentially upregulated by ZnSO_4_ treatment and did not include any associated with the ISR. Instead, they included early stress response transcription factors such as EGR1, FOS, FOSL or IER5 indicating different stress responses between Zn2+ and GSK-J4 treatment, despite an overlapping metal response pattern.

**Figure 5:**
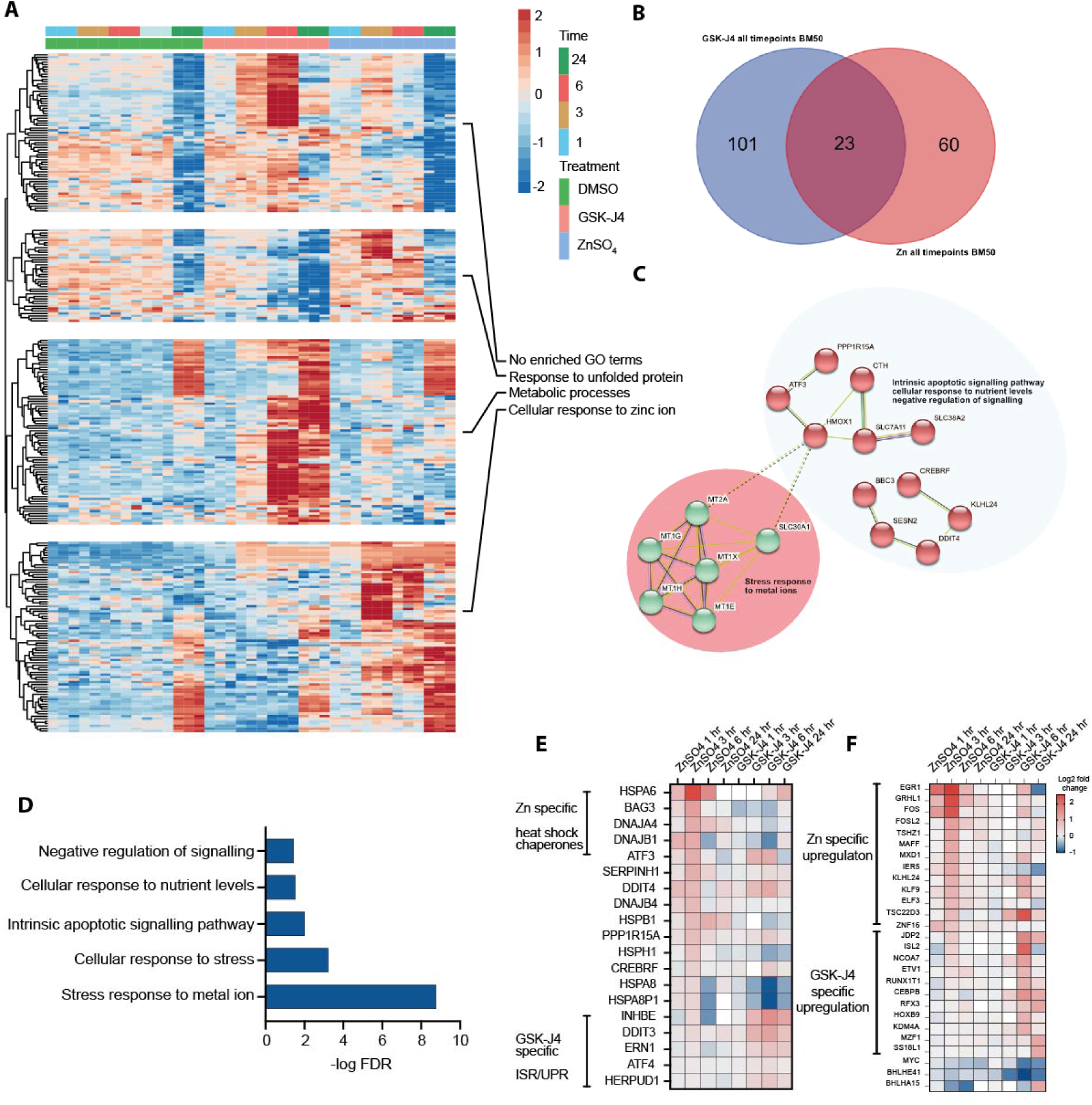
Comparison of JJN3 transcriptomes between GSK-J4 and ZnSO_4_ treatment. **(A)** Heatmap of all differentially expressed genes in JJN3 cells treated with either DMSO (0.1%), GSK-J4 (2 μM) or ZnSO_4_ (100 μM) after 1, 3, 6 and 24 hrs. **(B)** Venn diagram showing the overlap of all differentially regulated genes across all timepoints following treatment with GSK-J4 and ZnSO_4_. **(C)** STRING analysis of genes overlapping between GSK-J4 and ZnSO_4_. **(D)** GO analysis of showing the ontologies that are enriched for overlapping genes between GSK-J4 and ZnSO_4_. **(E-F)** Heatmaps showing the expression of selected stress response **(E)** or transcription and chromatin factor **(F)** genes following GSK-J4 or ZnSO_4_ treatment for 1, 3, 6 and 24 hrs.

## Conclusions

The transcriptional upregulation of metallothionein genes has been a hallmark of multiple studies using inhibitors of the histone demethylase KDM6 - observed in published RNA-seq datasets generated from different cell types as well as from primary myeloma cells and cell lines. We had originally hypothesized that KDM6 inhibition may directly impact the transcriptional response of the metallothionein locus by virtue of chromatin modification and subsequent gene regulation. However, our data from live-cell imaging and chelation assays demonstrate that GSK-J4 produces a transient surge in intracellular free Zn^2+^ concentrations. This suggests that direct metal chelation by GSK-J4 and subsequent transport of an inhibitor-Zn^2+^ complex across the plasma membrane is the source eliciting a transcriptional metal response signature. There is clear precedent of small molecules acting as ionophores, e.g. as observed for Zn^2+^ transmembrane transport mediated by chloroquine [42]. Accordingly, this increase in free intracellular Zn^2+^ provides a reasonable explanation for the increase in metallothionein gene expression that is observed in our study, however this does not rule out additional transcriptional mechanisms that are mediated through KDM6 inhibition. Although ZnSO_4_ treatment and KDM6 inhibition share a metallothionein response through MTF1, the two treatments diverge in other transcription factor repertoires they elicit. Previous studies have shown a hierarchy of Zn^2+^ response factors, with MTF1 as dominating transcription factor [41]. Differences in transcription factor repertoires, in particular the induction of the ATF4 mediated ISR, might explain the pro-apoptotic effects observed with KDM6 inhibition. Our previous studies in chordoma [22], NK cells [25], and Th17 cells [23] have consistently shown presence of a dual metallothionein and ATF4 driven response upon GSK-J4 treatment, where the latter may contribute to phenotypic responses such as cytokine regulation in immune cells. Available data suggest that GSK-J4 is not an unselective cellular toxic agent [13, 23]. While immune cells do not display signs of apoptosis but undergo a reversible proliferation block as observed with T-cells [23] certain cancer cell types such as chordoma and myeloma experience cell death upon KDM6 inhibition. In chordoma this effect is due to direct regulation of the essential oncogenic transcription factor brachyury (TBXT) by KDM6, resulting in increased repressive H3K27me3 marks thus silencing the oncogenic transcription factor [22]. Cancers such as myeloma have a high secretory protein output which requires complex mechanisms to control proteotoxic stress [43, 44]. The observed cell death upon KDM6 inhibition in myeloma cells may therefore be related to the ISR dominated by pro-apoptotic DDIT3 expression, an observation with possible therapeutic implications, justifying further studies.

## MATERIALS and METHODS

### Multiple Myeloma cell lines

The human multiple myeloma cell lines JJN3, RPMI-8226, MM1R, and NCI-H929 were gifts from Prof Claire Edwards, NDORMS, University of Oxford. The MM1S cell line was a gifts from Prof Gareth Morgan. AMO-1 and JJN3 cells lines were purchased from DSMZ (Braunschweig, Germany) or LGC standards (Teddington, UK). All multiple myeloma cell lines were cultured in RPMI (Sigma) with the addition of 10% heat inactivated Fetal Calf serum (Invitrogen), supplemented with 2mmol/L glutamine (Sigma). Cell lines were tested and authenticated by short tandem repeat DNA fingerprinting analysis (Public Health England culture collection, www.phe-culturecollections.org.uk). All cell lines were regularly tested for mycoplasma contamination by PCR.

### Cell viability

Cell viability was measured using resazurin-based PrestoBlue assay (ThermoFisher) according to the manufacturer’s instructions. Briefly, 10 μL of PrestoBlue was added to 100 μL of media containing cells and incubated for 30 minutes at 37°C. Plates were read on a FLUOstar Optima (BMG) with excitation filter at 535nm and emission filter at 615nm.

### Compound Screening

The epigenetic tool compound library was compiled from collaborator and commercial sources (SI DATA 1). Myeloma cells were plated at 5,000 cells per well in 96 well plates in a volume of 90 ml of medium. Compounds were dissolved at 1000x final concentration in DMSO, diluted to 10x final concentration in media and 10 ml of compound was added to each of the experimental wells. Each experiment was performed in triplicate. Cells were grown in the presence of compound for 5 days.

Cell viability was assessed using a resazurin oxidation assay by the addition of 10ul of sodium resazurin (Sigma) (0.01% in PBS) to each well, with incubation at 37°C for 2 hours followed by reading the fluorescence (Ex 560nm, Em590nm) in a plate reader. Results from compound treated wells are expressed as a percentage of the control wells present on each plate. For EC_50_ determination, cells were seeded at 10,000 cells/well into 96-well plates and treated with increasing concentrations of the test compound. Cell viability was detected as described after 5 days. Data were fitted to calculate EC_50_ values using Graphpad Prism.

### Flow cytometry

For flow cytometry detection, cells were harvested and washed once in PBS. Apoptosis was determined by detecting the presence of FITC-conjugated Annexin V using the Apoptosis Detection kit (Abcam) and cell death was measured using Propidium Iodide (PI) staining. Briefly, following compound treatment, cells were harvested, washed once in PBS and resuspended in 500 μL of binding buffer containing 5 μL of AnexinV-FITC and 5 μL of PI (50 ug/mL). Cells were incubated in the dark for 10 minutes before being analysed using a LSR Fortessa (BD) flow cytometry running FACSDiva Software version 6.

### Assessment of cellular proliferation by BrdU incorporation

Cellular proliferation was measured using the FITC BrdU Flow Kit (BD Pharminagen). Briefly, cells were treated with compound for 24 hours. BrdU was added at a final concentration of 10 µM and cells harvested after 30 minutes, 1 hour and 3 hours. Cells were fixed and permeabilized and stored at −80°C in 90%FCS, 10% DMSO. To stain, cells were thawed, washed and re-permeabilized prior to incubation with DNase 30 µg/ 10^6^ cells for 1 hour. FITC-labelled anti-BrdU antibody was added to cells for 20 min and total DNA stained with 7-aminoactinomycin D. Samples were kept in the dark prior to analysis.

### Microarray expression profiling

For microarray studies, RNA was isolated using Trizol reagent (Ambion) according to the manufacturer’s instructions. Resulting RNA was quantified and quality controlled using a Nanodrop spectrophotometer ND1000 (Thermo Fisher). cDNA was prepared using the ABI high capacity cDNA synthesis kit, (Invitrogen) including RNAsin in the reactions (Promega). Samples were analysed using the HumanHT12v4 Expression BeadChip Kit (Illumina). Data was processed using in R (version 3.1.1) using the “Bioconductor” and “Lumi” packages.

### Transcriptomic/Bulk RNA-seq

Myeloma cells at a density of 1×10^6/ml were treated for 6 and 24 hours with 1µM of GSK-J4 or DMSO-treated as controls. All conditions were performed in triplicate. Approximately 300,000 cells were harvested for each sample for RNA extraction. Cells were centrifuged for 5 minutes at 1500rpm, the supernatant was removed, and the cell pellets were resuspended in 300ul of Trizol. 6-hour samples were stored at −80°C overnight, and 24 hour samples were kept on ice prior to RNA extraction. RNA was extracted using Direct-zol RNA Miniprep kits (Zymo Research, USA), following the manufacturer’s instructions. RNA was quantified using a NanoDrop^TM^ 2000 spectrophotometer and standardised to 100ng of RNA in 50ml of nuclease free water. Library preparation was performed using Nebnext Ultra II directional RNA library prep kit for Illumina with Poly(A) mRNA Magnetic Isolation Module (New England Biolabs, USA), following the manufacturer’s instructions. Libraries were quantified using a 2200 Tapestation (Agilent Technologies, USA) with high sensitivity D1000 screen tapes. Libraries were denatured and diluted prior to sequencing. Paired-end sequencing was performed using the Illumina NextSeq 500, according to the manufacturer’s instructions.

### Bulk RNAseq analysis

FASTQ files were downloaded from the Basespace platform. A cgat-core pipeline within the cribbslab repository (‘pipeline_pseudobulk.py’; (https://github.com/cribbslab/cribbslab/blob/master/cribbslab/pipeline_pseudobulk.py) was used to process the reads [45]. Kallisto [46] was implemented for pseudo-alignment of the reads, with a K-mer size of 31 base pairs. Homo sapiens (human) genome assembly GRCh38 (hg38) transcriptome was used as a reference. Differential expression analysis was performed using DESeq2 [47]. Pathway analysis was performed using the XGR package [48] and gene ontology (GO) annotations [49].

### Reverse transcription quantitative PCR analysis

Following total RNA extraction and DNAseI digestion by using TRIzol reagent (Invitrogen) and a direct-zol RNA Miniprep kit (Zymo research), complementary DNA (cDNA) was generated by using a SuperScript II RT kit (Invitrogen), as per manufacturer’s instructions. Reverse transcription PCR was performed with specific primers to determine expression levels for each gene. Quantified mRNA was normalised to the relative expression of β-actin gene. All experiments were performed in technical triplicates. The primers used for MTX1 in this study are forward: 5’–TCTCCTTGCCTCGAAATG-3’, reverse: 5’–CACAGCTGTCCTGCCATC −3’.

### KDM6 inhibition using human BM samples followed by single-cell transcriptomics

Bone marrow samples were collected from two newly diagnosed multiple myeloma patients; anonymised human tissue samples used in this project were obtained with informed consent by the HaemBio Tissue Bank (REC reference: 17/SC/0572). After Ficoll gradient separation, mononuclear bone marrow cells were diluted to 500,000 cells/ml in RPMI media supplemented with 2 mM L-glutamine and 10% FBS, and 1 ml was added to 15 ml polypropylene tubes. Compounds were dissolved in DMSO, and 1 mL of compound solution was added to achieve a final concentration of 1 µM and incubated for 24 hours. Cells were counted, and single-cell RNA-seq library preparation was performed using the Chromium Next GEM Single Cell 3’ GEM, Library & Gel Bead Kit v3.1, according to the manufacturer’s instructions. Indexed libraries were quantitated by TapeStation, pooled, and sequenced on an Illumina NovaSeq 6000 (Novogene, UK).

### Single-cell RNAseq analysis

FASTQ files were downloaded from GEO using the accession number GSE186448 using the SRA-toolkit. We used scflow pipeline (https://github.com/cribbslab/scflow) to process 10X chromium scRNA-seq reads. The pipeline is a wrapper for the Kallisto BUS/ BUStools workflow [50] and was implemented to pseudo-align reads to a reference transcriptome. Homo sapiens (human) genome assembly GRCh38 (hg38) transcriptome was used as a reference. The output was converted to single-cell experiment object. Quality control and filtering was performed and then converted to a Seurat object for downstream analysis. Any cells with a mitochondrial ratio higher than 0.1, fewer than 500 UMIs, or fewer than 300 or greater than 6000 features were removed. Clustering and sample integration was performed using Seurat [51]. Automated cell type annotation was performed using singleR [52], clustifyr [53] and scClassify [54]. The celldex HumanPrimaryCellAtlasData reference annotation was used for singleR and clustifyr. Annotation with ScClassify was performed with the author’s pretrained model ‘Joint Human PBMC’ comprising seven annotated PBMC scRNA-seq data sets. Seurat’s Wilcoxon model was used for differential expression analysis.

### Fluorescence microscopy to measure free intracellular zinc

Cells were treated with Fluozin-3 AM (Life Technologies), a cell-permeable fluorescent dye with selectivity for zinc over other divalent cations. 5µM Fluozin-3 AM was added to cell media and incubated at 37°C for 60 minutes. Cells were washed 3 times with PBS and then incubated for a further 60 minutes prior to treatment with GSK-J4. Images were obtained every 2 minutes using a Zeiss Axio microscope and images analysed using ZEN Lite software (Zeiss)

### Statistical analysis

Experiments were performed independently at least 3 times, and biological triplicates were used in each experiment unless otherwise specified. Data were analyzed using Kruskal-Wallis test followed by Dunn’s multiple comparison within the Graphpad software (GraphPad Software 9.0.1, La Jolla, CA, USA). (not significant N.S.; \**P* < .05; \*\**P* < .01). Error bars represent standard deviation.

## ACKNOWLEDGMENTS

This work was supported through Innovate UK (UO, MP, APC), the National Institute for Health Research Oxford Biomedical Research Centre (JED, UO), Cancer Research UK (CRUK, UO and JC), the Bone Cancer Research Trust (APC and UO), the Leducq Epigenetics of Atherosclerosis Network (LEAN) program grant from the Leducq Foundation (UO), the Chan Zuckerberg Initiative (APC) and the Myeloma Single Cell Consortium (UO). APC is a recipient of an MRC Career Development Fellowship (MR/V010182/1). The work was supported by the Wellcome Trust (Multi-user equipment grant “The London Metallomic Facility” (202902/Z/16/Z) (WM). We thank our collaborators Nick LaThangue (Oxford), Jian Jin (Mt Sinai), Xiang Wang (U Colorado), Elisabeth Martinez (U Texas) and the Structural Genomics Consortium (Paul Brennan, Peter Brown, Stefan Knapp, Susanne Muller Knapp) for providing compounds to the epigenetic screen.

## Data availability

Microarray and RNA sequencing data can be obtained from the GEO database using accession number GSE60600 and GSE213294, respectively. ZnSO and GSK-J4 treated Multiple Myeloma cells lines can be accessed from the GEO database using the accession number GSE186448. Publicly available data from NK cells, Th17 cells, Neurobloastoma cells and Adenocarcinoma cell lines can be accessed under the GEO database accession numbers GSE156423, GSE127767, GSE110709, GSE133970, respectively.

## Notes

### Competing Interest Statement

A.P.C. M.P and U.O. are cofounders of Caeruleus Genomics Ltd and are inventors on several patents related to sequencing technologies filed by Oxford University Innovations. Other authors declare no competing interests.

## REFERENCES

1. Brown, K.H., et al., Effect of supplemental zinc on the growth and serum zinc concentrations of prepubertal children: a meta-analysis of randomized controlled trials. Am J Clin Nutr, 2002. 75(6): p. 1062–71.

2. Prasad, A.S., Zinc in human health: Effect of zinc on immune cells. Molecular Medicine, 2008. 14(5-6): p. 353–357.

3. Maret, W., Zinc in Cellular Regulation: The Nature and Significance of "Zinc Signals". Int J Mol Sci, 2017. 18(11).

4. Kagi, J.H. and B.L. Valee, Metallothionein: a cadmium- and zinc-containing protein from equine renal cortex. J Biol Chem, 1960. 235: p. 3460–5.

5. West, A.K., et al., Human metallothionein genes: structure of the functional locus at 16q13. Genomics, 1990. 8(3): p. 513–8.

6. Takahashi, S., Molecular functions of metallothionein and its role in hematological malignancies. J Hematol Oncol, 2012. 5: p. 41.

7. Hamer, D.H., Metallothionein. Annu Rev Biochem, 1986. 55: p. 913–51.

8. Xin, X., et al., MiR-376a-3p increases cell apoptosis in acute myeloid leukemia by targeting MT1X. Cancer Biol Ther, 2022. 23(1): p. 234–242.

9. Liu, T.Y., D.J. Li, and J.F. Zhou, Effect of Metabolic Reprogramming on the Biological Activity of Acute Myeloid Leukemia Cells. Blood, 2018. 132.

10. Krezel, A. and W. Maret, The Bioinorganic Chemistry of Mammalian Metallothioneins. Chem Rev, 2021. 121(23): p. 14594–14648.

11. Liu, Z., et al., Metallothionein 1 family profiling identifies MT1X as a tumor suppressor involved in the progression and metastastatic capacity of hepatocellular carcinoma. Mol Carcinog, 2018. 57(11): p. 1435–1444.

12. Jiang, H., P.J. Daniels, and G.K. Andrews, Putative zinc-sensing zinc fingers of metal-response element-binding transcription factor-1 stabilize a metal-dependent chromatin complex on the endogenous metallothionein-I promoter. J Biol Chem, 2003. 278(32): p. 30394–402.

13. Kruidenier, L., et al., A selective jumonji H3K27 demethylase inhibitor modulates the proinflammatory macrophage response. Nature, 2012. 488(7411): p. 404–8.

14. Zhao, S., C.D. Allis, and G.G. Wang, The language of chromatin modification in human cancers. Nat Rev Cancer, 2021. 21(7): p. 413–430.

15. Wang, Y., et al., Beyond the double helix: writing and reading the histone code. Novartis Found Symp, 2004. 259: p. 3–17; discussion 17-21, 163–9.

16. Allis, C.D., et al., New nomenclature for chromatin-modifying enzymes. Cell, 2007. 131(4): p. 633–6.

17. De Santa, F., et al., The histone H3 lysine-27 demethylase Jmjd3 links inflammation to inhibition of polycomb-mediated gene silencing. Cell, 2007. 130(6): p. 1083–94.

18. Boyer, L.A., et al., Polycomb complexes repress developmental regulators in murine embryonic stem cells. Nature, 2006. 441(7091): p. 349–53.

19. Lee, T.I., et al., Control of developmental regulators by Polycomb in human embryonic stem cells. Cell, 2006. 125(2): p. 301–13.

20. Sevinc, K., et al., BRD9-containing non-canonical BAF complex maintains somatic cell transcriptome and acts as a barrier to human reprogramming. Stem Cell Reports, 2022. 17(12): p. 2629–2642.

21. Cribbs, A.P., et al., Dissecting the Role of BET Bromodomain Proteins BRD2 and BRD4 in Human NK Cell Function. Front Immunol, 2021. 12: p. 626255.

22. Cottone, L., et al., Inhibition of Histone H3K27 Demethylases Inactivates Brachyury (TBXT) and Promotes Chordoma Cell Death. Cancer Res, 2020. 80(20): p. 4540–4551.

23. Cribbs, A.P., et al., Histone H3K27me3 demethylases regulate human Th17 cell development and effector functions by impacting on metabolism. Proc Natl Acad Sci U S A, 2020. 117(11): p. 6056–6066.

24. Ebrahimi, A., et al., Bromodomain inhibition of the coactivators CBP/EP300 facilitate cellular reprogramming. Nat Chem Biol, 2019. 15(5): p. 519–528.

25. Cribbs, A., et al., Inhibition of histone H3K27 demethylases selectively modulates inflammatory phenotypes of natural killer cells. J Biol Chem, 2018. 293(7): p. 2422–2437.

26. Loven, J., et al., Selective inhibition of tumor oncogenes by disruption of super-enhancers. Cell, 2013. 153(2): p. 320–34.

27. Santo, L., et al., Preclinical activity, pharmacodynamic, and pharmacokinetic properties of a selective HDAC6 inhibitor, ACY-1215, in combination with bortezomib in multiple myeloma. Blood, 2012. 119(11): p. 2579–89.

28. Mithraprabhu, S., et al., Histone deacetylase (HDAC) inhibitors as single agents induce multiple myeloma cell death principally through the inhibition of class I HDAC. Br J Haematol, 2013. 162(4): p. 559–62.

29. Gulla, A., et al., Protein arginine methyltransferase 5 has prognostic relevance and is a druggable target in multiple myeloma. Leukemia, 2018. 32(4): p. 996–1002.

30. Kurata, K., et al., Prolyl-tRNA synthetase as a novel therapeutic target in multiple myeloma. Blood Cancer J, 2023. 13(1): p. 12.

31. Nowak, R.P., et al., Advances and challenges in understanding histone demethylase biology. Curr Opin Chem Biol, 2016. 33: p. 151–9.

32. Johansson, C., et al., Structural analysis of human KDM5B guides histone demethylase inhibitor development. Nat Chem Biol, 2016. 12(7): p. 539–45.

33. Chu, X., et al., GSK-J4 induces cell cycle arrest and apoptosis via ER stress and the synergism between GSK-J4 and decitabine in acute myeloid leukemia KG-1a cells. Cancer Cell Int, 2020. 20: p. 209.

34. Das, A., et al., RNA sequencing reveals resistance of TLR4 ligand-activated microglial cells to inflammation mediated by the selective jumonji H3K27 demethylase inhibitor. Sci Rep, 2017. 7(1): p. 6554.

35. Zhang, Z.M., et al., SAPCD2 promotes neuroblastoma progression by altering the subcellular distribution of E2F7. Cell Death Dis, 2022. 13(2): p. 174.

36. Lochmann, T.L., et al., Targeted inhibition of histone H3K27 demethylation is effective in high-risk neuroblastoma. Sci Transl Med, 2018. 10(441).

37. Subramanian Vignesh, K. and G.S. Deepe, Jr., Metallothioneins: Emerging Modulators in Immunity and Infection. Int J Mol Sci, 2017. 18(10).

38. Cho, Y.E., et al., Cellular Zn depletion by metal ion chelators (TPEN, DTPA and chelex resin) and its application to osteoblastic MC3T3-E1 cells. Nutr Res Pract, 2007. 1(1): p. 29–35.

39. Choi, J.S., et al., Inhibition of cyclooxygenase-2 expression by zinc-chelator in retinal ischemia. Vision Res, 2006. 46(17): p. 2721–7.

40. Gee, K.R., et al., Detection and imaging of zinc secretion from pancreatic beta-cells using a new fluorescent zinc indicator. J Am Chem Soc, 2002. 124(5): p. 776–8.

41. Hardyman, J.E., et al., Zinc sensing by metal-responsive transcription factor 1 (MTF1) controls metallothionein and ZnT1 expression to buffer the sensitivity of the transcriptome response to zinc. Metallomics, 2016. 8(3): p. 337–43.

42. Xue, J., et al., Chloroquine is a zinc ionophore. PLoS One, 2014. 9(10): p. e109180.

43. Moscvin, M., M. Ho, and G. Bianchi, Overcoming drug resistance by targeting protein homeostasis in multiple myeloma. Cancer Drug Resist, 2021. 4: p. 1028–1046.

44. Adams, J., The development of proteasome inhibitors as anticancer drugs. Cancer Cell, 2004. 5(5): p. 417–21.

45. Cribbs, A., et al., CGAT-core: a python framework for building scalable, reproducible computational biology workflows [version 2; peer review: 1 approved, 1 approved with reservations]. F1000Research, 2019. 8(377).

46. Bray, N.L., et al., Near-optimal probabilistic RNA-seq quantification. Nat Biotechnol, 2016. 34(5): p. 525–7.

47. Love, M.I., W. Huber, and S. Anders, Moderated estimation of fold change and dispersion for RNA-seq data with DESeq2. Genome Biol, 2014. 15(12): p. 550.

48. Fang, H., et al., XGR software for enhanced interpretation of genomic summary data, illustrated by application to immunological traits. Genome Med, 2016. 8(1): p. 129.

49. Harris, M.A., et al., The Gene Ontology (GO) database and informatics resource. Nucleic Acids Res, 2004. 32(Database issue): p. D258–61.

50. Melsted, P., V. Ntranos, and L. Pachter, The barcode, UMI, set format and BUStools. Bioinformatics, 2019. 35(21): p. 4472–4473.

51. Satija, R., et al., Spatial reconstruction of single-cell gene expression data. Nat Biotechnol, 2015. 33(5): p. 495–502.

52. Aran, D., et al., Reference-based analysis of lung single-cell sequencing reveals a transitional profibrotic macrophage. Nat Immunol, 2019. 20(2): p. 163–172.

53. Fu, R., et al., clustifyr: an R package for automated single-cell RNA sequencing cluster classification. F1000Res, 2020. 9: p. 223.

54. Lin, Y., et al., scClassify: sample size estimation and multiscale classification of cells using single and multiple reference. Mol Syst Biol, 2020. 16(6): p. e9389.

