## Supplementary Figures for "The KDM6 histone demethylase inhibitor GSK-J4 induces metal and stress responses in multiple myeloma cells"

**Supplementary Figure 1**

**GSJ-K4 and GSK-J5 dose response curves in various Multiple Myeloma cell lines.**

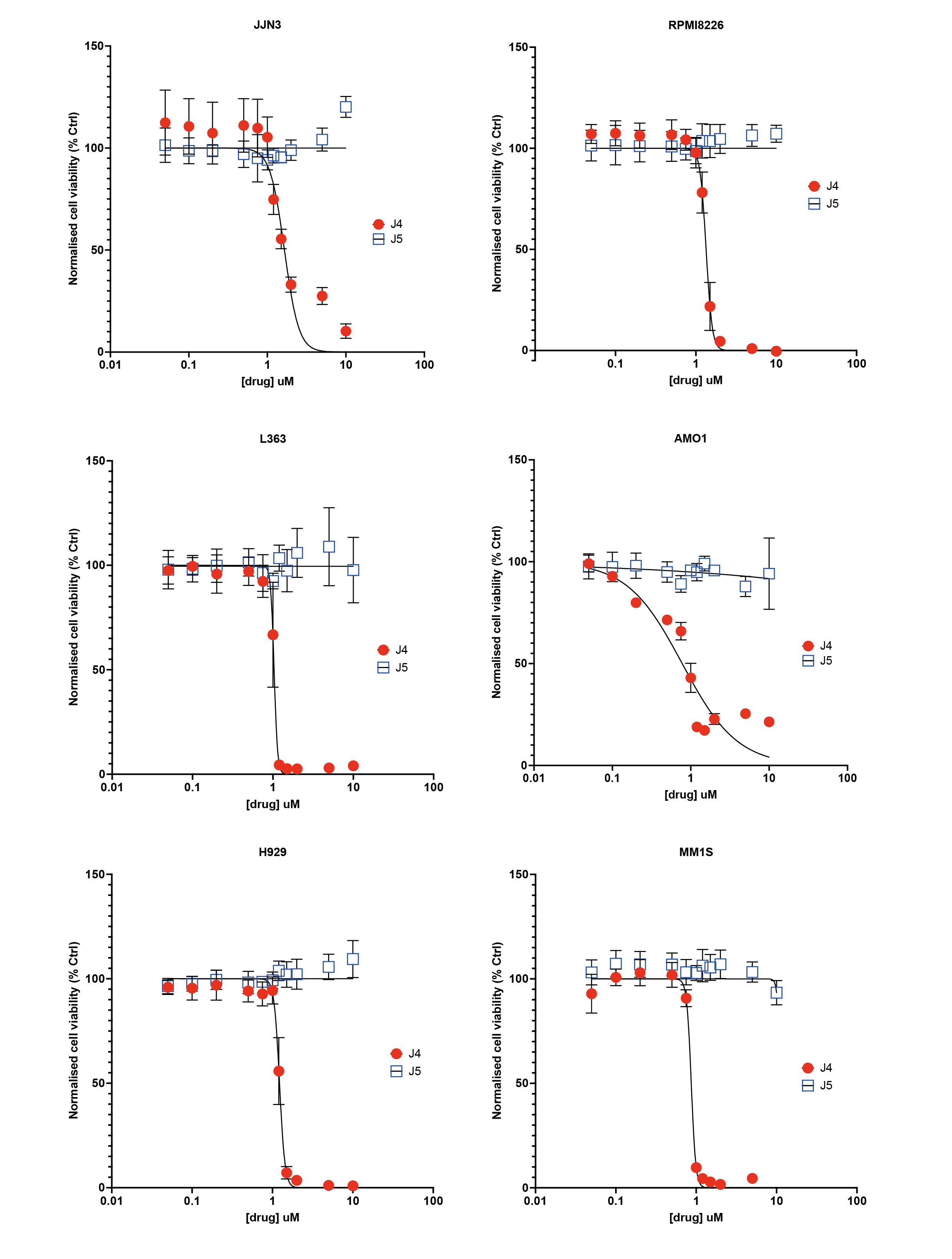

**Supplementary Figure 2**

**Antiproliferative effects of GSK-J4 measured by EDU incorporation.**

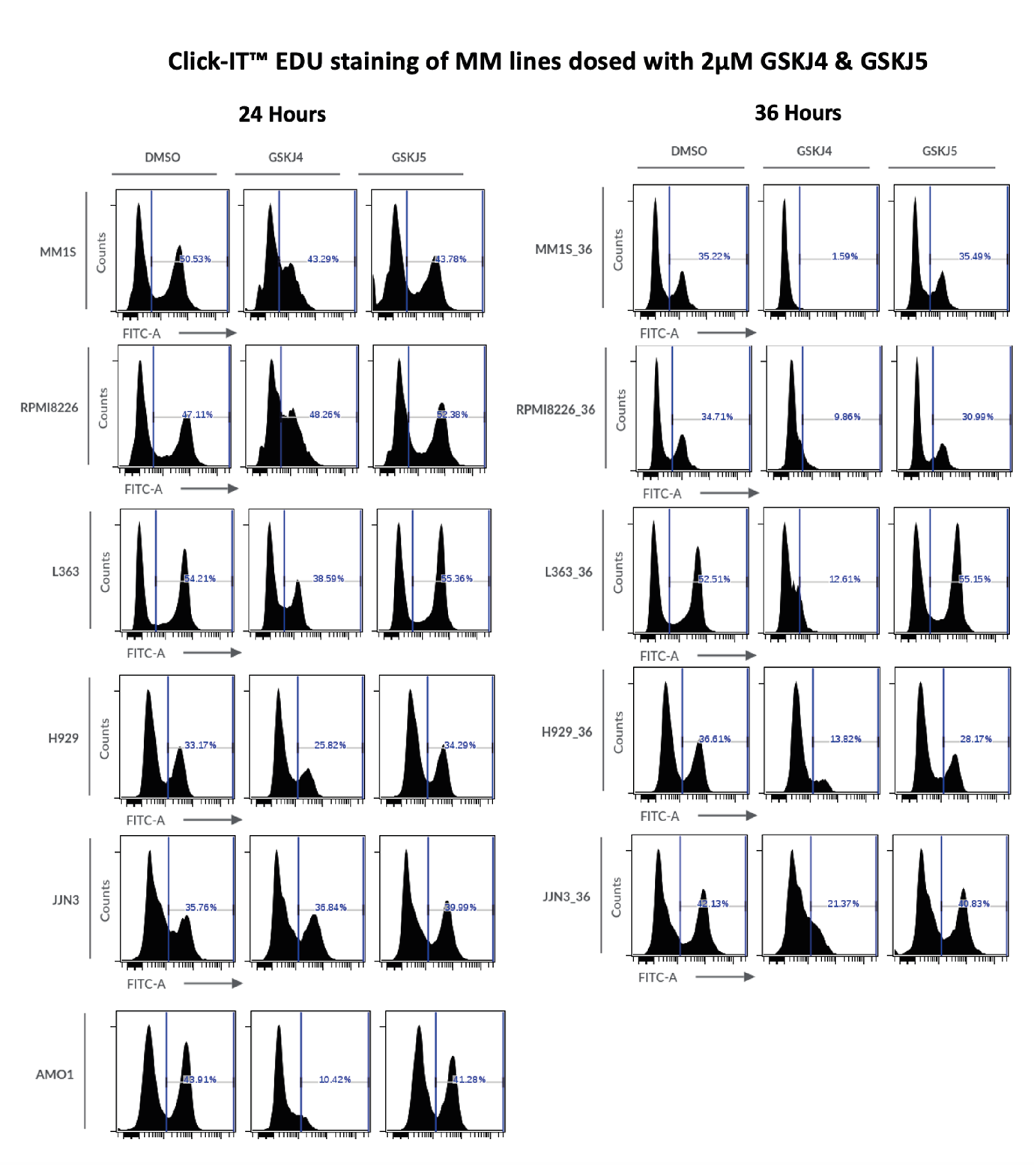

**Supplementary Figure 3**

**Cell-death effects of GSK-J4 measured by Annexin V and PI staining.**

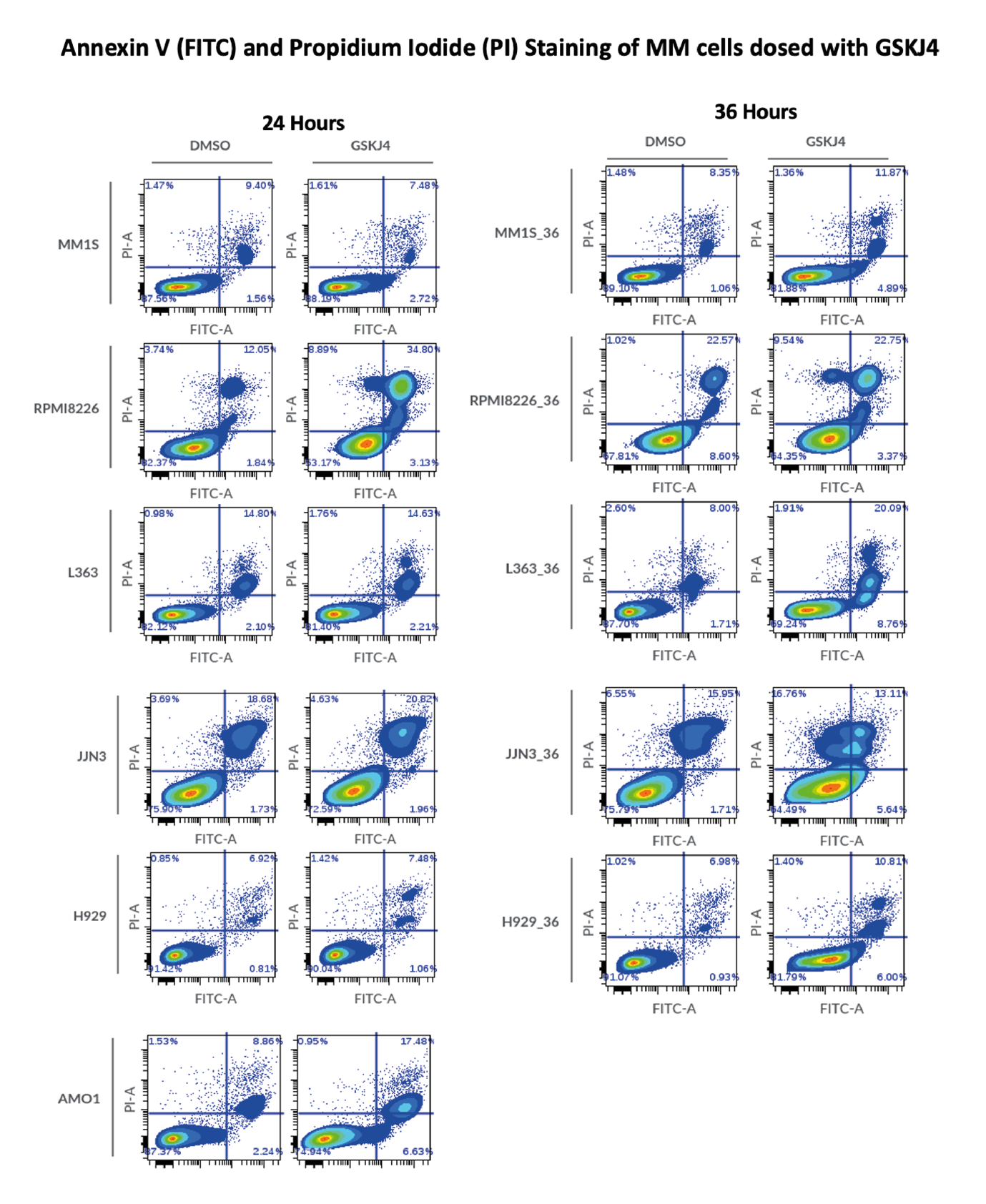

**Supplementary Figure 4**

**Annexin V (FITC) and Propidium Iodide (PI) Staining of MM cells dosed with GSKJ4 for 72hrs**
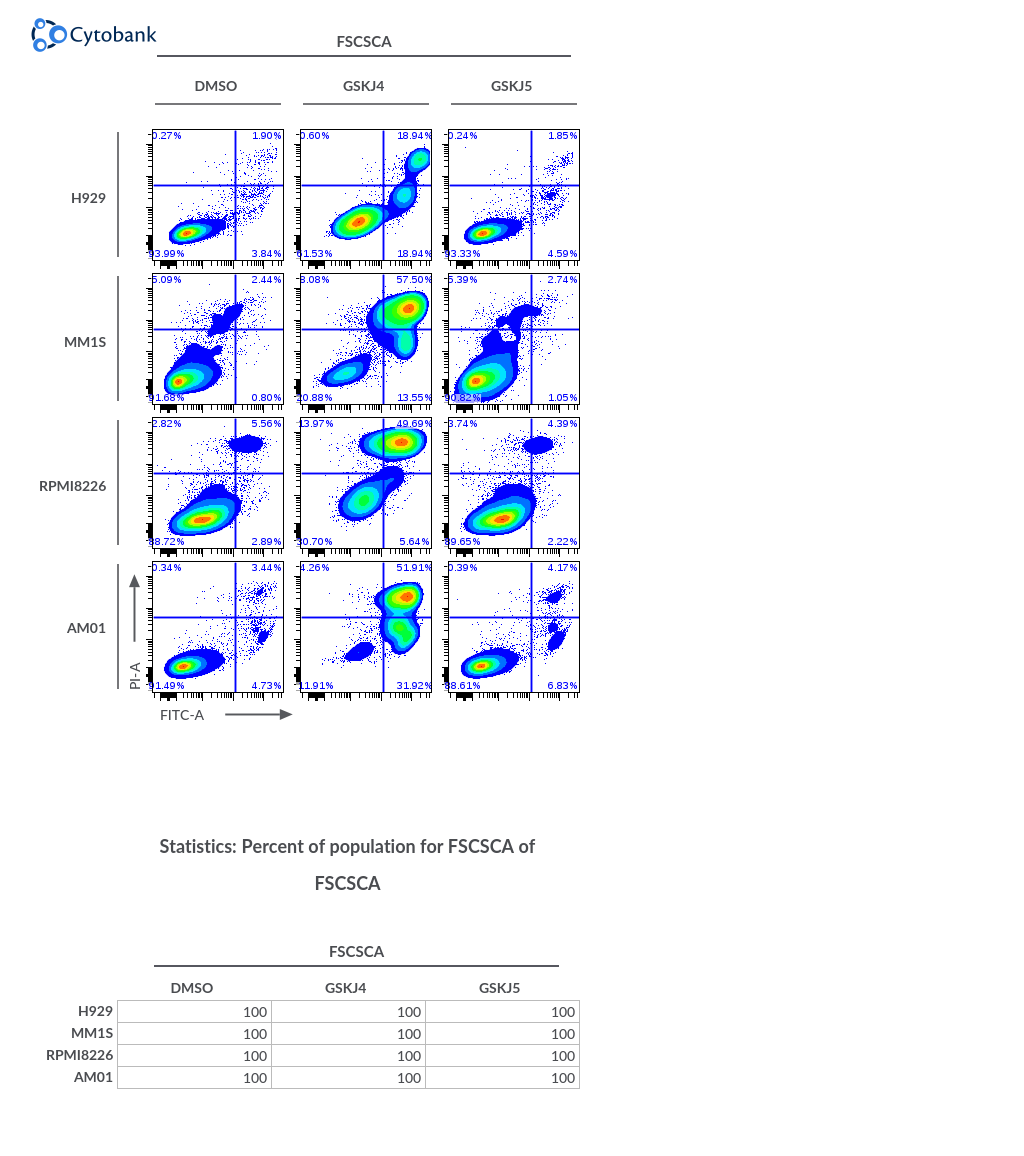

**Supplementary Figure 5**

Heatmap depicting metallothionein and SLC expression levels (RNAseq data) upon GSK-J4 treatment in different cellular systems.

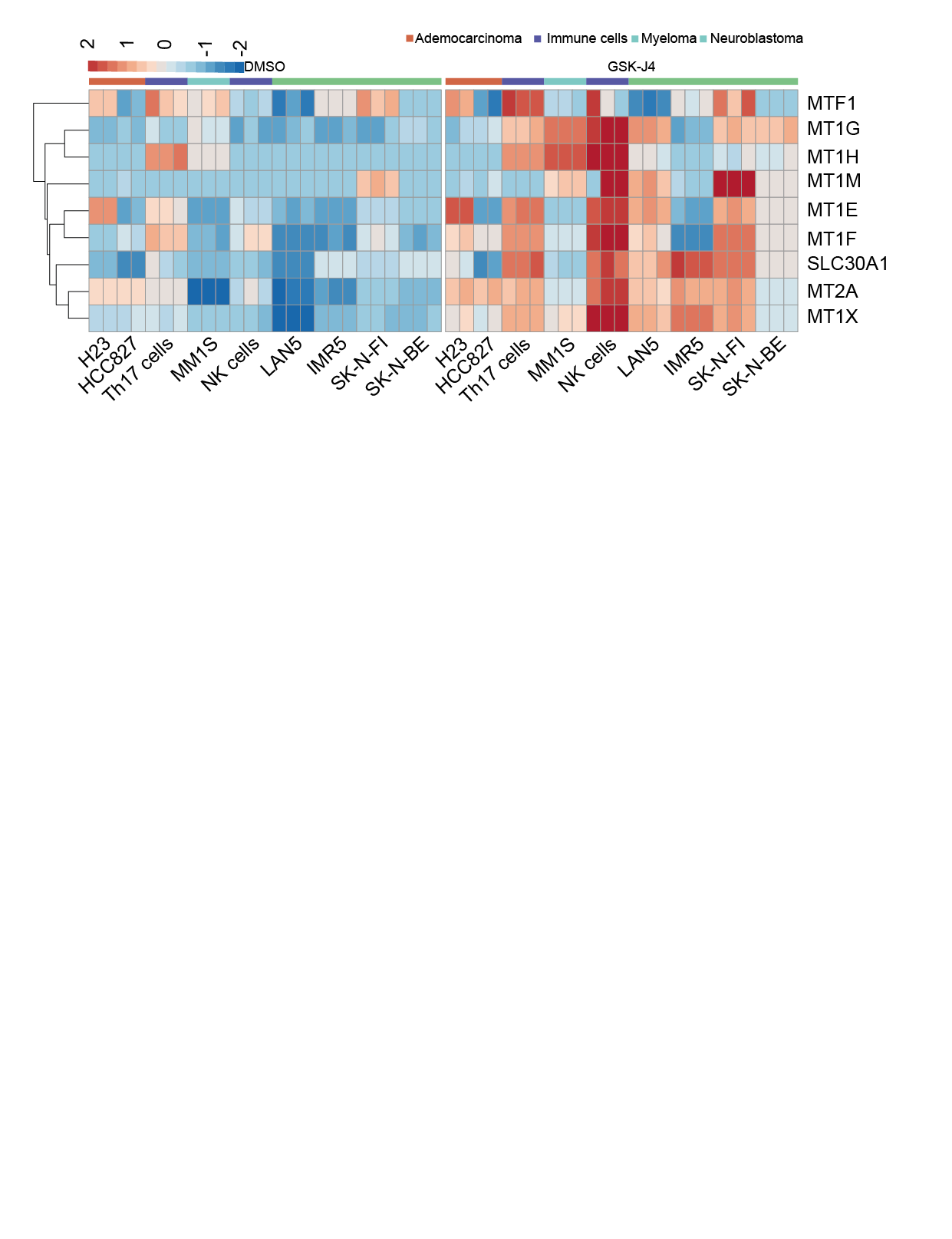

**Supplementary Figure 6**

UMAP showing the distribution of cell types in a Myeloma patient sample. UMAP plots showing the expression of MT1X (left panels) and MT1G (right panels) treated with either DMSO or GSK-J4.

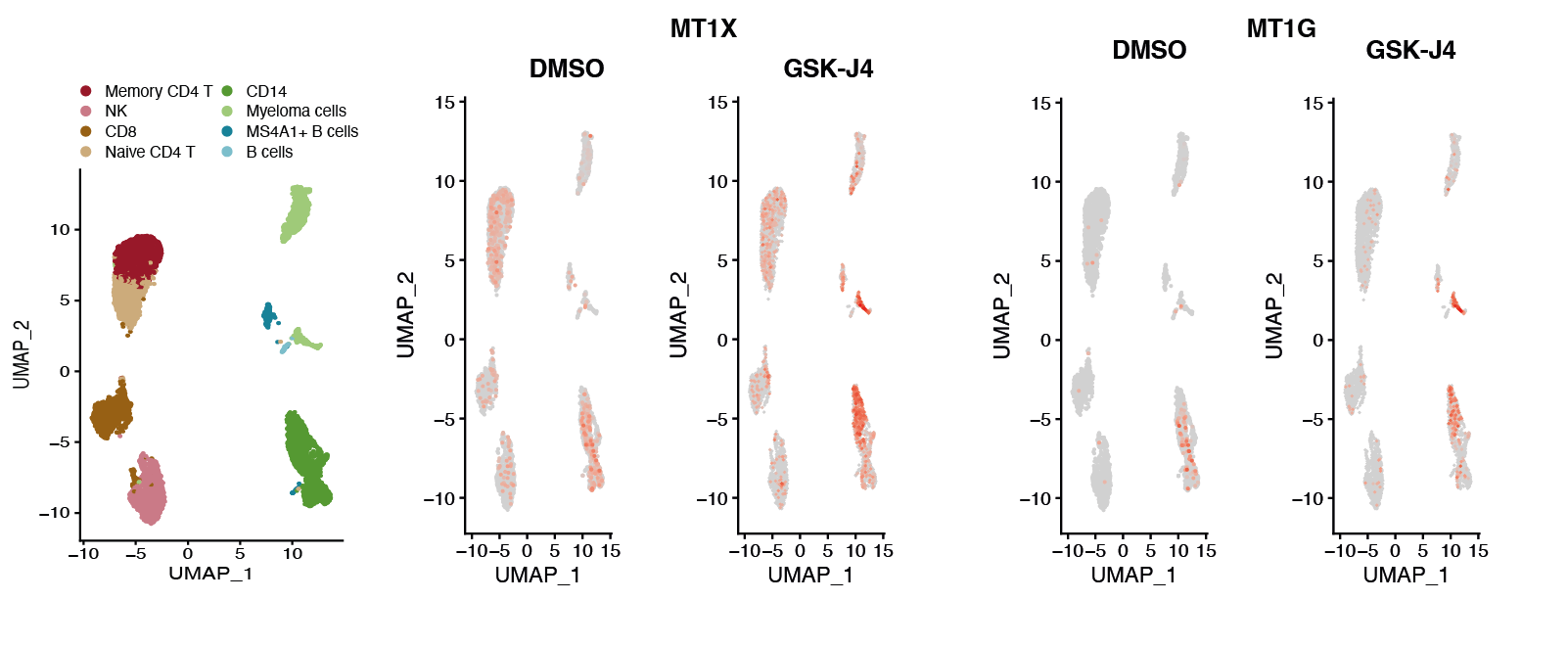
